# Improved breeding for *Fusarium pseudograminarium* (Fusarium crown rot) using qPCR measurement of infection in multi-species winter cereal experiments

**DOI:** 10.1101/2022.10.12.512005

**Authors:** Andrew Milgate, Brad Baxter, Steven Simpfendorfer, Nannan Yang, Beverly Orchard, Ben Ovenden

## Abstract

Fusarium crown rot (FCR) causes significant grain yield loss in winter cereals around the world. Breeding for resistance and/or tolerance to FCR has been slow with relatively limited success. In this study, multi-species experiments were used to demonstrate an improved method to quantify FCR infection levels at plant maturity using qPCR, as well as the genotype yield retention using residual regression deviation. Using qPCR to measure FCR infection allowed a higher degree of resolution between genotypes than traditional visual stem basal browning assessments. The results were consistent across three environments with different levels of disease expression. The improved measure of FCR infection along with genotype yield retention allows for partitioning of both tolerance and partial resistance. Together these methods offer new insights to FCR partial resistance and its relative importance to tolerance in bread wheat and barley. This new approach offers a more robust, cost-effective way to select for both FCR traits within breeding programs.

**Key message:** Genetic gain for tolerance and partial resistance against Fusarium crown rot (FCR) in winter cereals has been impeded by laborious and variable visual measures of infection severity. This paper presents results of an improved method to quantify FCR infection that are strongly correlated to yield loss and reveal previously unrecognised partial resistance in barley and wheat varieties.

## Introduction

Fusarium crown rot (FCR) is an important disease of winter cereals in many cereal growing regions around the world (Smiley and Patterson, 1996; Gargouri et al., 2011; Zhang et al., 2015; Alahmad et al., 2018; Kazan and Gardiner, 2018). Economic impacts of FCR, primarily through reduced grain yields, are estimated to cost the Australian winter cereals industry (wheat and barley) AUD$97 million per annum (Murray and Brennan, 2009). The disease is caused by several *Fusarium* species which colonise the basal vascular tissues of tillers and restrict both water and nutrient translocation within the plant during grain fill (Knight and Sutherland, 2016). Of the *Fusarium* species that can be the causal agent of FCR, *F. pseudograminarium* (*Fp*) is the most common species observed in Australia, whereas *F. culmorum* is found to be associated with the disease in cooler regions (Backhouse and Burgess, 2002; Akinsanmi et al., 2004; Backhouse et al., 2004; Poole et al., 2013). However, both *Fusarium* species are known to often co-exist in the same locations.

FCR infections occur frequently in conservation cropping systems where tight rotations of cereal crops and retention of cereal residue (stubble) are practised (Simpfendorfer et al., 2019). Stubble acts as a refuge for the pathogen to survive up to three years or more. *Fusarium* mycelia colonises stubble residue, which then provides the inoculum source for infecting the next cereal crop or alternate hosts, including grasses such as *Phalaris, Agropyron* and *Bromus* species (Purss, 1969; Summerell and Burgess, 1988; Summerell et al., 1989; Summerell et al., 1990; Burgess, 2014). Soil inoculum levels, symptom development and grain yield losses are influenced by environmental factors and the types of cereals being grown (Hollaway et al., 2013). FCR infection can occur early in seedling development right through to adult stages of growth. When moisture stress occurs during flowering and grain-filling, infected tillers can senesce prematurely to appear as ‘white heads’. These white heads are absent of grain or contain few shrivelled grains. FCR can impact several components of grain yield including kernel number per head, kernel weight, tiller height and straw weight (Smiley et al., 2005). FCR infection occurs in both susceptible and partially resistant genotypes (Percy et al., 2012; Knight and Sutherland, 2015; 2016). FCR inoculum levels and impacts on grain yield are significant only in years with rainfall below the long-term average during the grain-filling period and minimised in relatively wet years (Hollaway et al., 2013). Hence, an alternative for FCR resistance breeding is to conjoin FCR partial resistance and/or tolerance with stress related physiological traits such as drought, heat or moisture stress (Kazan and Gardiner, 2018). While these studies have not given definitive explanations of the molecular basis for the associations, they suggest that drought stress related responses are beneficial in the presence of FCR infection (Su et al., 2021).

The goal of breeding programs is to achieve high grain yield but with stable performance in the presence of disease. Breeders of winter cereals have had great success in protecting yields to *Puccinia* species and *Zymoseptoria tritici* around the world by combining major and minor resistance genes in resistance breeding programs (Raman and Milgate, 2012; Ellis et al., 2014; Brown et al., 2015). Breeding for FCR resistance currently focuses on identifying genotypes with partial resistance that reduces the development of basal stem browning symptom. Studies in segregating crosses or association panels have identified QTLs for resistance on at least 13 of the 21 wheat chromosomes (Collard et al., 2006; Ma et al., 2010; Liu and Ogbonnaya, 2015; Martin et al., 2015; Rahman et al., 2020; Rahman et al., 2021). However, these QTLs fail to be widely utilised in breeding programs because of the complex nature of inheritance, partial effectiveness, and poor agronomic performance including low yield potential (Rahman et al., 2020). Therefore, for FCR there is little effective resistance deployed, and all varieties are infected and suffer losses to some extent (Kazan and Gardiner, 2018). Tolerance, the ability to retain yield in the presence of infection, is frequently discussed in studies examining genotype effects of FCR and how it differs from partial resistance (Kazan and Gardiner, 2018; Forknall et al., 2019). Differences in tolerance are known to exist between winter cereal species. Barley is rated more tolerant than bread wheat which is more tolerant than durum wheat (Hollaway et al., 2013). However, difficulties exist to experimentally distinguish clear phenotypic differences between tolerance and partial resistance in field studies. Improved methods for selecting more resistant varieties are urgently needed because there are currently no effective available fungicide control options that consistently prevent yield losses from FCR.

Typically, detailed yield assessments at multiple levels of disease intensity are required to measure FCR tolerance accurately (Forknall et al., 2019). Disease assessments are performed either by visual scoring of stem basal browning lesions or more recently by using qPCR to measure pathogen DNA load in seedlings (Liu et al., 2012; Knight and Sutherland, 2015; Ozdemir et al., 2020). Both methods have large environmental interactions, are labour intensive and have low heritability (Dodman and Wildermuth, 1987; Knight and Sutherland, 2015; Martin et al., 2015; Kazan and Gardiner, 2018; Rahman et al., 2020; Kelly et al., 2021) and are not compatible with the selection of genotypes with combinations of multiple genes with minor effects within large breeding populations (Rahman et al., 2021).

In this study, we use multi-species experiments to estimate the genotype yield potential in treatments with (inoculated) and without (non-inoculated) FCR infection in three target environments, across two years. FCR infection was measured at maturity using qPCR, which allowed a quantitative measurement of *Fp* infection loads to identify genotypes with partial resistance. Increased infection levels were strongly correlated with grain yield loss. Partitioning of tolerance and partial resistance of genotypes was achieved by using residual regression deviation as described in Kelly et al. (2021). This study is the first to measure *Fp* DNA at plant maturity in the field as a proxy for FCR infection severity and associate it with yield loss. Previous reports of using measures of pathogen DNA have only been conducted under controlled environmental conditions in seedlings and have not associated measures with yield loss from FCR (Liu et al., 2012; Ozdemir et al., 2020).

## Materials and methods

### Plant materials

Sixteen wheat, eight barley and one durum wheat genotypes were used to conduct these field experiments (Table 1). These genotypes reflected the mix of commercial cultivars grown in NSW, Australia at the time the experiments where conducted. Each genotype has a known resistance rating to FCR, which ranged from moderately susceptible (MS) to very susceptible (VS). All genotypes were included in all experiments, with the exception of one barley variety, Buloke, which was present only in the Wagga Wagga 2016 field experiment.

**Table 1.**
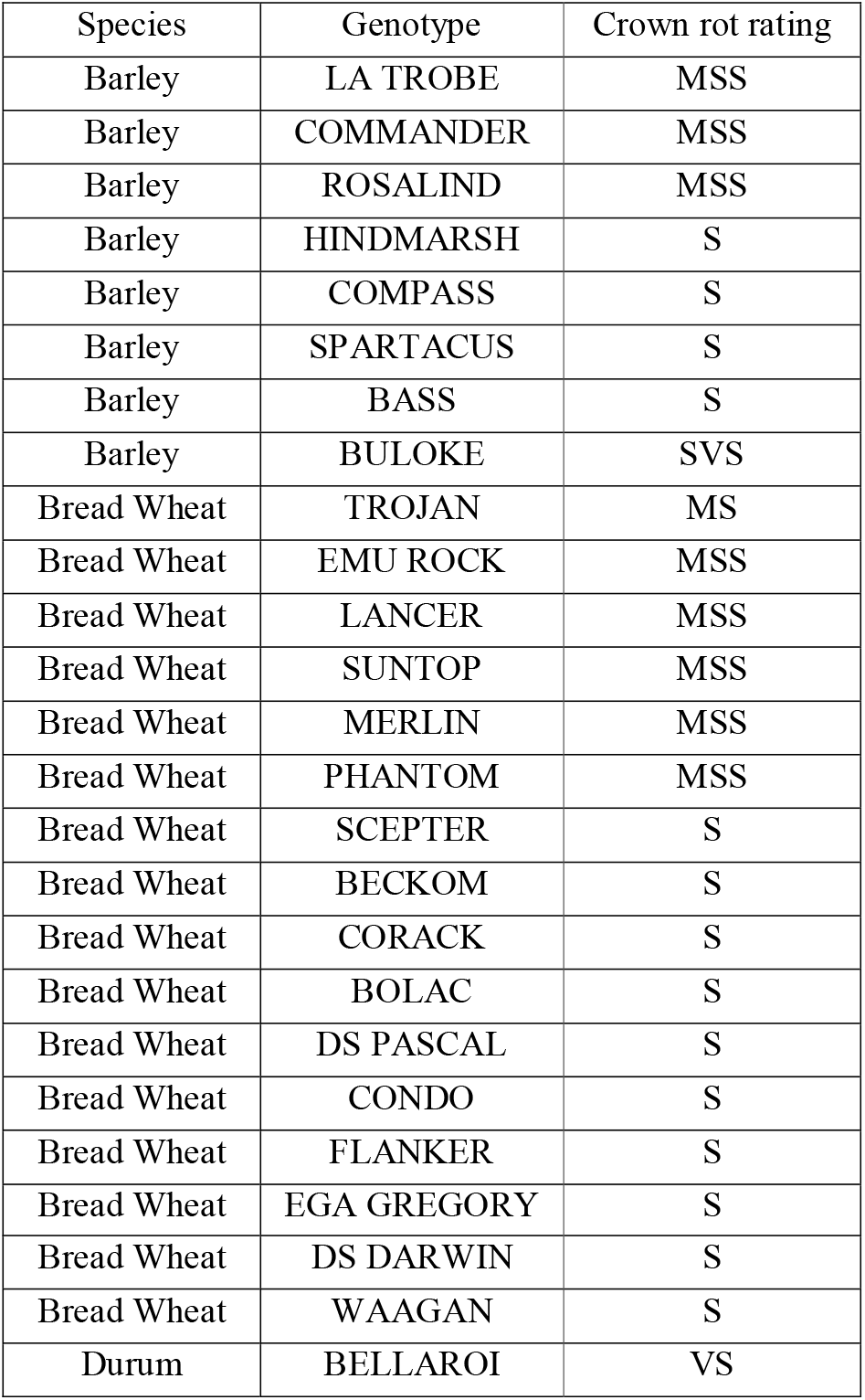
Barley, bread wheat and durum wheat genotypes included in the experiments had known crown rot resistance ratings (Crown rot rating source Winter crop variety sowing guide 2015-2017).

### Experimental design

The package DiGGer (Coombes, 2002) in R (R Core Team, 2021) was used to create randomised complete block (spatial) designs for all experiments in the study. Treatments with inoculum (inoculated) and no inoculum (non-inoculated) were randomised, each with four replicates in independent experiments.

### Field experiments

The genotypes were evaluated in three different environments, over two years. The site by year combinations and rainfall totals including annual, in-crop growing season (June to November) and during grain-filling (September or October) are outlined in Table 2. Field experiments were conducted at Wagga Wagga Agricultural Institute at Wagga Wagga (S-35.04419222, E147.3167896), NSW, Australia between 2016-2017 and the Condobolin Agricultural Research and Advisory Station at Condobolin (S-33.064939, E147.230877), NSW, Australia in 2017. Experiments were managed with standard agronomic practices for the region. Foliar diseases were prevented from impacting on grain yield by targeted in-crop fungicide applications at key growth stages. Seed rates at sowing were calculated using 1,000 grain weight and germination percentage to target the optimal number of plants per metre square (m^2^) for each site based on regional practice.

**Table 2:** Rainfall statistics for each experimental site. Growing season (GS) rainfall June-November Wagga Wagga, June-October Condobolin, is from a sowing to maturity in each trial. Grain fill is during October at Wagga Wagga and September for Condobolin. ^#^Note: Mean rainfall figures calculated separately for each site from weather statistics available at the CSIRO Bureau of Meteorology; Wagga Wagga AMO 1941-2021 and Condobolin Agricultural Research Station 1954-2022.

| Location | Year | Annual rainfall | Decile | GS rainfall (June-Nov) | Grain fill rainfall | Mean Grain fill rainfall<sup>#</sup> | Mean GS rainfall<sup>#</sup> |
| --- | --- | --- | --- | --- | --- | --- | --- |
| Wagga Wagga | 2016 | 778.8 | 9 | 496 | 64 | 56 | 308 |
| Wagga Wagga | 2017 | 445.3 | 2 | 215 | 65 | 56 | 308 |
| Condobolin | 2017 | 490.6 | 6 | 104 | 6 | 32 | 180 |

Sowing date was 2-4 weeks after the optimal sowing window for each experimental site. This coincided with a 1st week of June sowing date compared to the commercial best practice window of late April to early May. This increased the probability and extent of moisture and heat stress during grain filling, further exacerbating the expression of FCR. The experimental plot inoculation was carried out at sowing, according to the grain inoculum method described by Forknall et al. (2019) and Kadkol et al. (2021). Briefly, two grams of inoculum per metre of trial row was homogenised with the viable seed in packets and sown with a plot seeder. Viable seed only was sown in the non-inoculated treatment.

Disease severity was measured at harvest in all replicates of both the inoculated and non-inoculated treatments in each of the field experiment using quantitative PCR (qPCR). The sampling methodology ensured that the residual stubble on the outside rows and ends of the plots were avoided. A random collection of stubble from the mature plants after grain harvest, with an even coverage across internal rows of the plots was undertaken. The stubble was trimmed to a uniform length of 5 cm, ensuring the crown and first node was present with 32 random pieces per plot homogenised into 500 grams of pre-sterilised soil prior to analysis. The soil was sterilised twice on alternate days to remove any background levels of *Fusarium* species in an autoclave at 121°C and 115kpa for 60 minutes. The samples were then sent to South Australian Research and Development Institute (SARDI) for qPCR analysis (Ophel-Keller et al., 2008). Three tests were selected from the qPCR panel to be undertaken on the samples including, *F. pseudograminearum* (*Fp*) test 1 and test 2, and a third test which detects both *F. culmorum/graminearum*. The *F. culmorum/graminearum* test was included to detect any background FCR infection by the main non-target *Fusarium* spp., no significant infections were observed, only 5 plots out of 592 detected DNA of *F. culmorum/graminearum* above trace levels. For analysis the sum of *Fp* test 1 and test 2 were used as the surrogate for total FCR infection severity.

### Phenotypic data analysis

Phenotype records for the four traits (grain yield non-inoculated (t/ha), grain yield inoculated (t/ha), disease severity non-inoculated (Log_10_ *Fp* pgDNA/gram), disease severity inoculated (Log_10_ *Fp* pgDNA/gram) modelled using a multivariate multiplicative mixed linear model using approach of (Gilmour et al., 1997). Records for two traits (disease severity noninoculated, disease severity inoculated) were log transformed before modelling. The model used is described as follows:

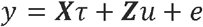

where y is a vector of length *n* = 4 × 296 containing stacked vectors for the four traits. *τ* is a vector of fixed effects including trait means for the design matrix *X*. The term *u* is the vector of genotype effects for each trait corresponding to the experimental design structure *Z*. The vector *e* of length *n* containing the residuals of the four traits was modelled with an unstructured variance-covariance matrix between traits. This structure permits the fitting of linear relationships at the residual level between the four traits. Modelling was performed using the R software package ASReml-R version 4 (Butler et al., 2017), in the R statistical software environment (R Core Team, 2021).

The linear relationships between each of the four traits was determined from the trait: genotype covariance modelling using methods detailed in Zhang et al. (2019) as follows:

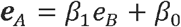

Where *A* and *B* refer to the two traits in each respective pairwise comparison between the four traits modelled, and where the slope of the regression is calculated as:

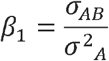

and the intercept *ß_0_* was determined from the overall genotype Best Linear Unbiased Predictions (BLUPs) for the four traits across experiments from the multivariate mixed model. The difference (residual) between the BLUP from the mixed model for the response trait in each pairwise comparison and the predicted value on the trend line was calculated. The deviation from the regression values are referred to hereafter as retention values.

## Results

Seasonal conditions varied considerably across the three experimental sites, resulting in differences across sites and treatments for yield (Table 2 & 3). Rainfall was above average during the growing season (GS) at Wagga Wagga in 2016. Below average GS rainfall was received at Wagga Wagga in 2017 and Condobolin in 2017, reaching only 70% and 57% of their annual mean rainfall respectively (Table 2). The range in moisture conditions allowed for differences in expression of tolerance and partial resistance to FCR infection to occur. FCR infection occurred in all experiments, with variation occurring in levels of infection between genotypes and treatments (Table 3). Low levels of infection were detected in all non-inoculated treatments across the three field experiments. Significant differences (*p* < 0.0001) of FCR infection severity (qPCR) were observed between treatments at all three sites (Table 3). The qPCR results discriminated the infection levels between genotypes in the inoculated treatments and non-inoculated treatments (Supplementary Table 1).

**Table 3:** Summary table of mean treatment effects for grain yield (t/ha) and disease severity of crown rot infection (Log_10_ pgDNA/gram) at each experiment. Non-inoculated = plots sown without crown rot inoculum, Inoculated = plots sown with crown rot inoculum. Inoculum was *Fusarium pseudograminarium*. Means for all experiments with p-value for paired comparison of treatments is given at the experiment level.

| EXPT | Treatment | N | CR infection ($\text{Log}_{10}$ pgDNA/gram) | p value | Grain yield (t/ha) | p value |
| --- | --- | --- | --- | --- | --- | --- |
| 2017 Condobolin | Non-inoculated | 100 | 2.17 | <0.0001 | 1.305 | 0.0027557 |
| 2017 Condobolin | Inoculated | 100 | 4.51 |  | 0.964 |  |
| 2017 Wagga Wagga | Non-inoculated | 100 | 0.27 | <0.0001 | 3.891 | 0.0006582 |
| 2017 Wagga Wagga | Inoculated | 100 | 3.79 |  | 3.511 |  |
| 2016 Wagga Wagga | Non-inoculated | 96 | 0.52 | <0.0001 | 5.379 | 0.4514391 |
| 2016 Wagga Wagga | Inoculated | 96 | 3.40 |  | 5.208 |  |

### Yield retention and partial resistance

Grain yield was reduced by FCR infection in the three field experiments and significant for Wagga Wagga in 2017 (*p* = 0.0006582) and Condobolin in 2017 (*p =* 0.0027557) (Table 3). Yield at the three sites was in accordance with the seasonal rainfall patterns (Table 2). The lowest site mean yield (1.305 t/ha) in the non-inoculated treatment was recorded at Condobolin in 2017 whilst the highest (5.379 t/ha) was measured at Wagga Wagga in 2016.

Significant variations in FCR infection severity were detected among genotypes using Log scores of the amount of *Fp* DNA per gram of stubble (Log_10_ *Fp* pgDNA/gram, Figures 1 – 3). Overall, the levels of qPCR FCR infection severity were 100-1000 times higher in the inoculated treatment than in the non-inoculated treatment (Table 3). The highest natural background infection severity in the non-inoculated treatment were measured in the barley genotype Bass (Log_10_ 1.215), while the highest background level detected in bread wheat was in the genotype Waagan (Log_10_ 1.163) (Supplementary Table 1). Both Bass (Log_10_ 4.119) and Waagan (Log_10_ 4.115) also had the highest infection severity in the inoculated treatment of all genotypes (Figure 1B, Supplementary Table 1). There was a strong positive correlation (*r* = 0.715) between the levels of FCR severity recorded for all genotypes in the two treatments (Figure 1B). Six out of eight barley genotypes had lower levels of FCR severity than the bread wheat genotypes included in the experiments.

**Figure 1.**
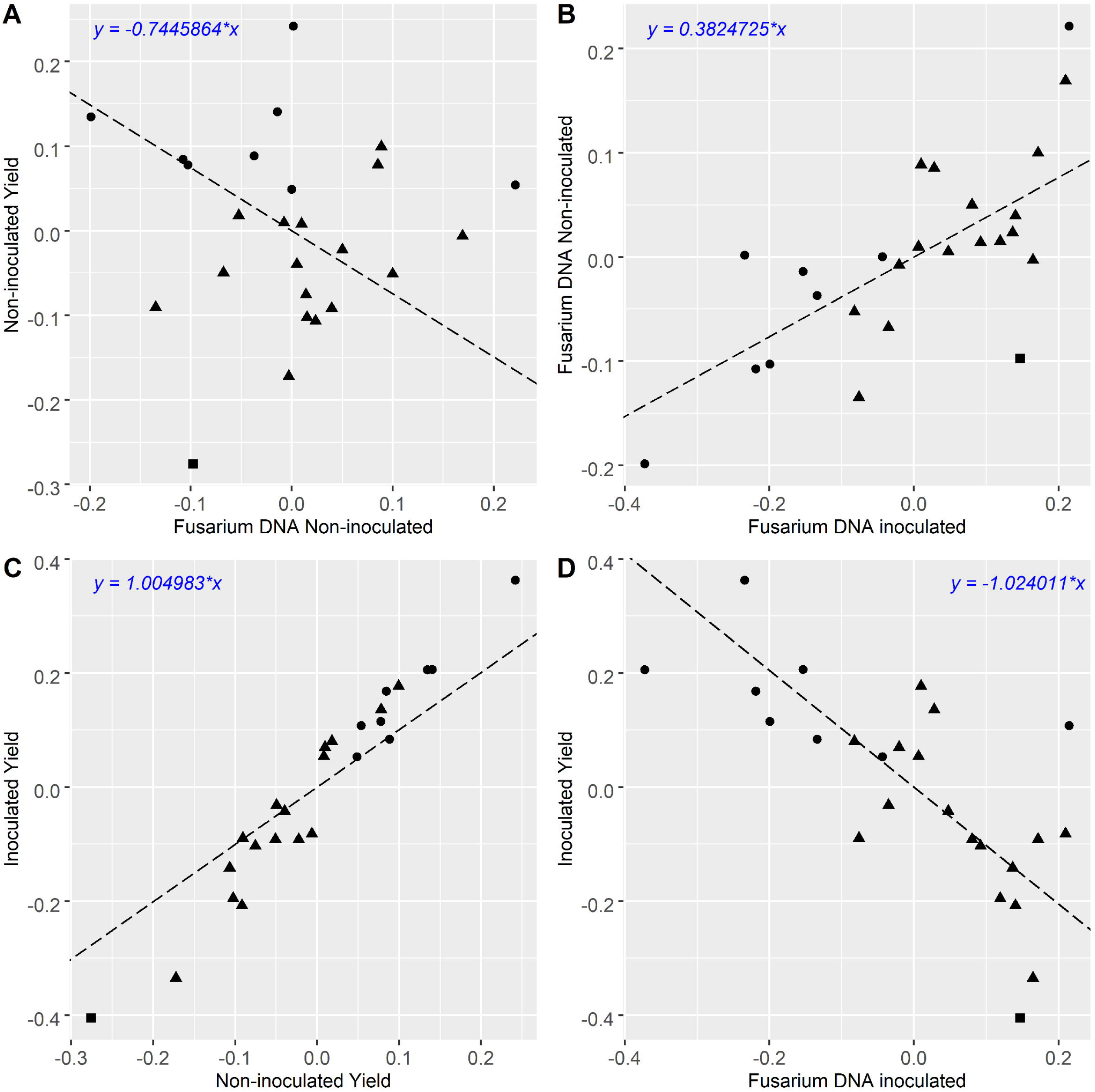
A-D: Comparison of grain yield and qPCR CR infection BLUPs of winter cereals from 3 experiments. Grain yield in t/ha, *Fusarium* DNA is qPCR of *F. pseudograminarium* in Log_10_ pgDNA/gram. Triangle, square and circle symbols indicate bread wheat, durum wheat and barley genotypes respectively. Pearson’s correlation coefficient for each comparison is A: *r* = 0.018, B: *r* = 0.715, C: *r* = 0.974, D: *r* = −0.715.

**Figure 2:**
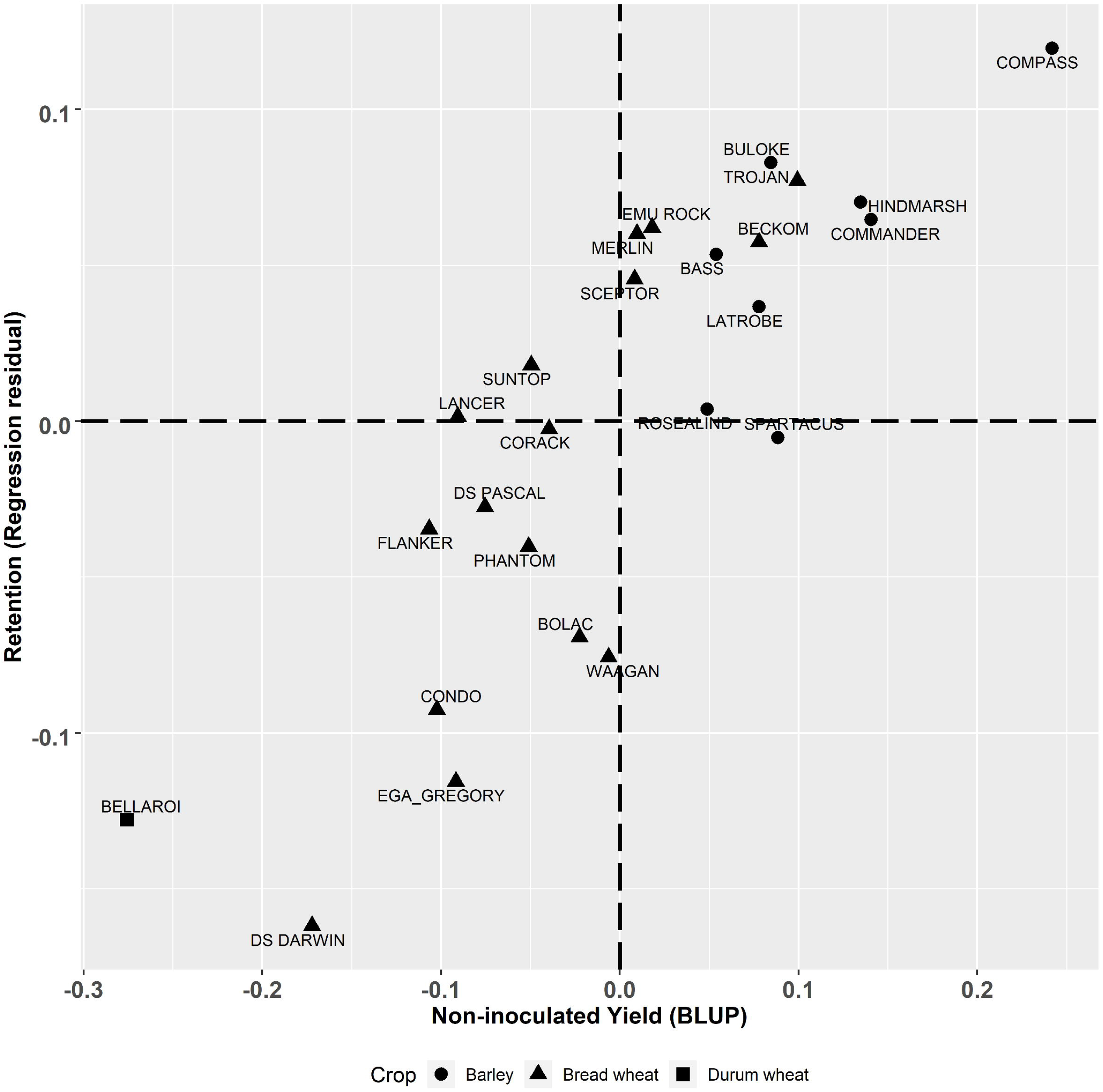
Best linear unbiased predictors (BLUPs) of genotype effects from the multivariate analysis for the derived trait of yield retention (t/ha) (regression of residuals) plotted against yield (t/ha) in non-inoculated plots. Triangle, square and circle symbols indicate bread wheat, durum wheat and barley genotypes respectively.

**Figure 3:**
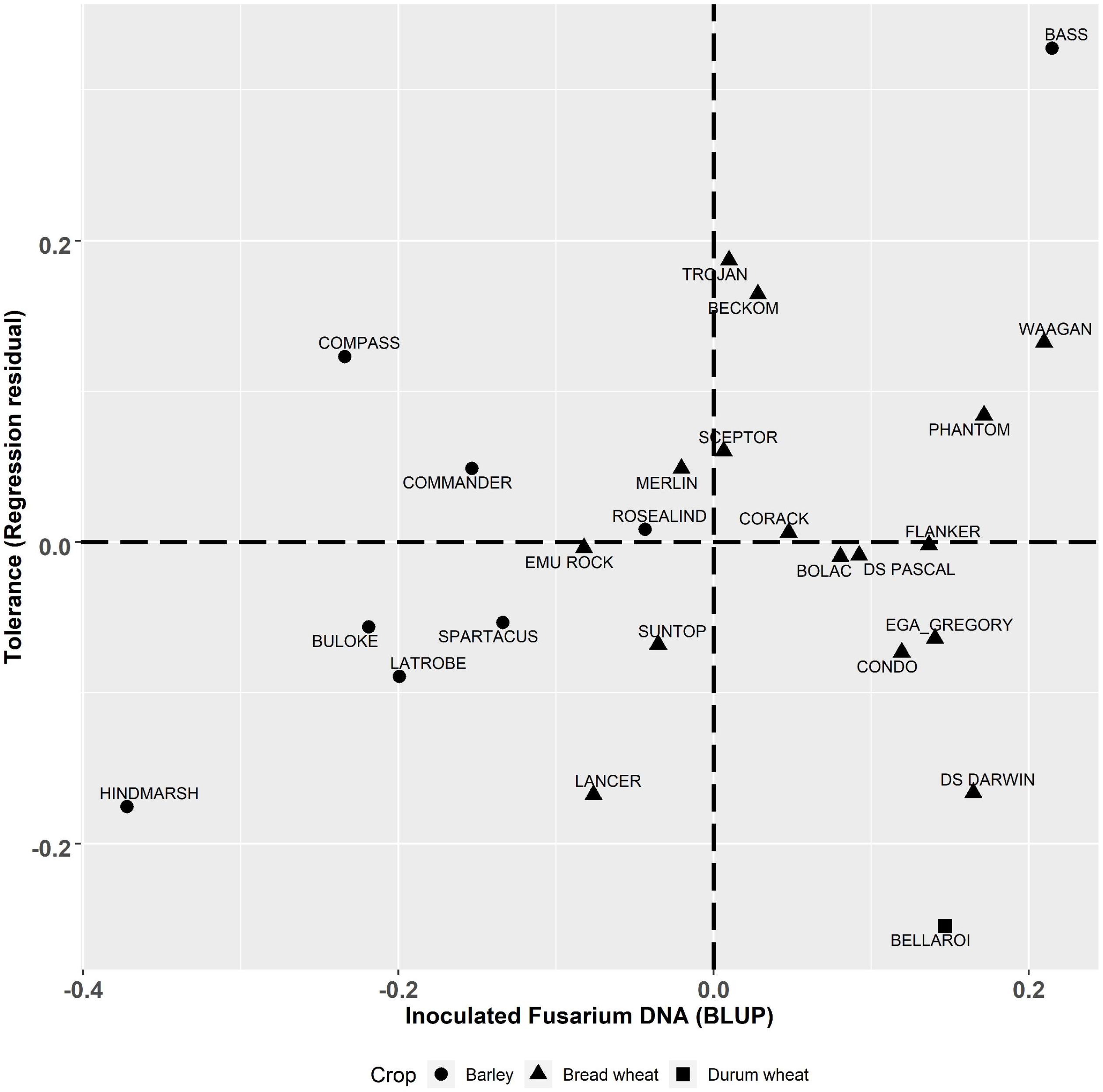
Best linear unbiased predictors (BLUPs) of genotype effects from the multivariate analysis for the derived trait of tolerance (t/ha) (regression residual) graphed against CR infection (Log_10_ pgDNA/gram) in the inoculated plots. Triangle, square and circle symbols indicate bread wheat, durum wheat and barley genotypes respectively.

The multivariate analysis of yield under two contrasting levels of disease had a high correlation of *r* = 0.974 (Figure 1C). Barley genotypes had higher yield compared to most of the bread wheat genotypes in both the inoculated and non-inoculated treatments at each site. Within the inoculated treatment, when comparing the same cereal species, there was large genotype differences in achieved yield. When compared to the combined experimental yield mean (Supplementary Table 1), mean bread wheat yield ranged from −0.335 to +0.177 t/ha of the combined mean whilst mean barley yield ranged from +0.108 to +0.362 t/ha. The highest yielding barley genotype in both treatments was Compass with LRBP Trojan being the highest yielding bread wheat genotype. The durum wheat genotype Bellaroi was the lowest yielding of all cereal species and genotypes.

The retention value of genotypes (residual deviation from the regression between the yield under inoculated and non-inoculated conditions, Figure 1C) is plotted against non-inoculated yield in Figure 2. Genotypes in the top left quadrant of Figure 2 have above average yield in both the inoculated and non-inoculated treatments. While those in the bottom left quadrant have above average yield in the non-inoculated treatment but lower than average yield in the inoculated treatment. Genotypes in the top right quadrant have below average yield in the non-inoculated treatment but above average yield in the inoculated treatment. While those in the bottom right have below average yield in both the inoculated and non-inoculated treatments. The magnitude of the retention value detected amongst the experimental genotypes ranged from −0.16 to +0.12 t/ha. Direct comparisons can be made between genotypes which show large differences in their ability to retain yield in the presence of FCR infection, such as Waagan and Scepter. The yield of Waagan is negatively impacted under high levels of FCR infection compared to Scepter which maintained above average yield in the *Fp* inoculated treatment (Figure 2).

The degree of partial resistance, measured as reduced severity using qPCR, within the tested genotypes was negatively correlated (*r* = –0.715) to yield (Figure 1D). However, some genotypes deviate significantly from the regression indicating varying levels of tolerance to FCR. The tolerance value is presented (residual deviation of genotype yield performance in the *Fp* inoculated treatment) against FCR severity (*Fp* pgDNA/gram) is presented in Figure 3. Genotypes in the top right quadrant have above average FCR severity (more susceptible) but have higher yield relative to other genotypes with the same level of disease severity, as such, showing higher tolerance to FCR. While those in the bottom right have above average FCR severity (more susceptible) but lower than average yield in the inoculated treatment, displaying intolerance to FCR. Genotypes in the top left have below average FCR severity (more partial resistance) in the *Fp* inoculated treatment and also above average yield (higher tolerance). While those in the bottom left have below average FCR severity (more partial resistance) in the inoculated treatment but lower than average yield in the *Fp* inoculated treatment, displaying intolerance to FCR.

This study highlights the complexity of selecting for genetic improvement for FCR resistance and tolerance in winter cereals. Of the genotypes included in the study, LRBP Trojan has the highest level of resistance according to traditional phenotyping methods with a MS rating (Table 1). However, LRBP Trojan performed poorly, being worse than five other bread wheat genotypes when using qPCR to measure FCR disease severity. Conversely, LRBP Trojan performed comparatively well for the tolerance measure, ranking highest of the bread wheat genotypes, illustrating it maintains yield in the presence of FCR infection through a tolerance mechanism rather than partial resistance. Using retention alone fails to identify the different ways cereal genotypes can achieve yield stability in the presence of FCR infection.

Another example, where using qPCR to quantify disease severity reveals different responses to FCR, can be seen in barley genotypes Hindmarsh and Commander. Both genotypes rank highly for yield retention (Figure 2). However, when their partial resistance and tolerance are separated (Figure 3), they display contrasting adaptations to FCR infection. Hindmarsh in this study demonstrated the highest level of partial resistance of all barley genotypes, but it was the second least tolerant genotype to FCR infection. Commander combines some partial resistance with some level of tolerance to achieve the highest yield retention rank of the barley genotypes examined in this study.

## Discussion

This study has demonstrated in multi-cereal species experiments, an improved method to estimate the genotype yield potential in the presence of FCR infection by using the residual regression deviation as a measure of yield retention in combination with qPCR analysis of *Fp* levels at harvest as a measure of partial resistance. The use of qPCR to measure FCR severity in cereal stubble at maturity, allowed a higher degree of resolution between genotypes than traditional visual browning assessments of stem bases. The results were consistent across three experiments under different environmental conditions which resulted in varying levels of FCR severity and yield impact. The improved measure of FCR severity allows for the partitioning and selection for both tolerance and partial resistance within winter cereal genotypes. Together these methods offer new insights into FCR partial resistance and its relative importance to tolerance in bread wheat and barley. It also provides a more robust technique to select for both traits within breeding programs which requires reduced labour and specialist skills compared with visual assessments which are more subjective and variable between multiple operators.

qPCR quantification of *Fusarium* DNA at maturity, as a measure of FCR severity, improved the correlation between infection and yield loss. This study is the first to measure *Fp* DNA at plant maturity under field conditions as a proxy for FCR severity and associate it with yield loss. Previous studies using measures of *Fusarium* DNA have been conducted under controlled environment conditions in seedlings and have not been then associated to yield outcomes. Two studies, Ozdemir et al. (2020) and Liu et al. (2012) applied qPCR and real time PCR to inoculated seedlings which were up to 7 weeks old. They found *Fusarium* DNA levels in seedlings were correlated to visual disease scores of seedlings for some wheat varieties, however there were inconsistencies observed where symptoms did not match the high or low levels of DNA detected. This supports our finding that traditional rating methods of genotypes based on visual scoring of stem basal browning symptoms is not a reliable predictor of infection levels, when compared to qPCR quantification. In contrast to our study Liu et al. (2012) observed in seedlings that barley genotypes were more susceptible to infection than the bread wheat entries used. However, multiple studies have found barley has lower yield loss to FCR compared to bread wheat and this attribute has been associated with higher levels of tolerance (e.g. Hollaway et al. 2013). This study reveals for the first time that barley varieties have higher levels of partial resistance at maturity than bread wheat, as well as higher tolerance, with both traits making contributions to increased yield performance in the presence of FCR infection.

The severity of FCR infection determined by qPCR in this study had a strong negative correlation with yield in the presence of disease. Traditional visual methods of measuring FCR severity on the basis of browning of stem bases have not been found to be strongly correlated with yield outcomes (Kelly et al., 2021; Rahman et al., 2021). The suggested reasons are the complex environmental influence on FCR symptom expression and the extent of yield loss suffered by winter cereals (Hollaway et al., 2013). These traditional visual methods also only provide assessment of partial resistance with no measure of FCR tolerance (Forknall et al., 2019). Our method provides a quantitative measurement of phenotype that can be used to identify differences in partial resistance more accurately which is less subjective than previous visual measurements. Partial resistance is determined by the reduction of *Fp* infection severity, measured as reduced colonisation of plant tissue objectively assessed using qPCR.

Our improved measure of partial resistance also improves the estimate of tolerance to FCR. This allows for selection of both partial resistance and tolerance traits with greater confidence. Following the method of Kelly et al. (2021), this study has been able to effectively separate partial resistance and tolerance effects contributing to yield retention in the presence of FCR infection. Estimation of tolerance relies on a precise and repeatable measure of disease severity so that differences in yield at a given level of pathogen burden, reflect adaptations other than resistance, leading to reduced yield loss. The correlation between disease severity (stem basal browning) and yield was lower (*r* = −0.19) in Kelly et al. (2021) compared to this study (qPCR) at (*r* = —0.715). Rahman et al. (2021) also found that the visual assessment of basal browning had low heritability and a poor correlation with yield in the presence of FCR infection. In this study, the correlation between yield in different genotypes in inoculated and non-inoculated treatments was very strong (r = 0.974) as in Kelly et al. (2021) (*r* = 0.95), which is notable. These findings suggest the yield effects of FCR infection on genotypes are relatively stable across environments and it is the visible disease severity symptoms which are variable due to environmental interactions.

This study of multi-cereal species shows there is genetic potential to improve yield performance of wheat in the presence of FCR infection. The results confirm that barley genotypes have better yield performance in the presence of disease, and this is not solely due to adaptive traits such as shorter growing season providing escape or tolerance to disease expression, but also involves higher levels of partial resistance. Hollaway et al. (2013) showed barley yield loss was less than bread wheat and durum wheat under FCR disease pressure. The findings were not equated to measurement of FCR severity nor did it associate disease symptoms with yield loss. Not all barley genotypes had superior yield over bread wheat genotypes in the presence of FCR infection, suggesting this is potentially an adaptive trait. Using traditional disease assessment methods such as stem basal browning, none of the barley varieties have high levels of partial resistance. The most resistant barley genotype in this study is rated as moderately susceptible to susceptible (MSS) (Table 1). However, measuring infection severity using qPCR did reveal barley genotypes with lower levels of disease than bread wheat genotypes.

Kelly et al. (2021) suggests that in isolation there is limited value of retention (responsiveness, or regression residual) as a breeding trait because it does not partition the tolerance response independently from partial resistance. While in theory the pursuit of partitioning these effects is desirable, the practical application of the method at the scales required by breeding programs to screen large segregating populations makes it impractical given the resources required to apply current visual disease phenotyping methods. Yield retention as a first screen for large populations, is a simpler, lower-cost approach than current more labour-intensive phenotyping methods. Yield retention alone will identify genotypes that have either a combination of both traits or high levels of either partial resistance or tolerance, but ultimately for a grower high yield in the presence of FCR infection is the overriding breeding objective. For example, the contrast between the barley genotypes Commander and Hindmarsh in our study illustrate this point, both have high yield retention (Supplementary Table 1). However, these two genotypes achieved this in different ways. Hindmarsh achieved higher yield through having low tolerance and higher partial resistance compared to Commander, which had higher tolerance but less resistance to FCR infection (Figure 3). Hence, the ability to partition and quantify these two independent traits is important from a breeding perspective but could be completed in a sequential manner to make greater genetic gain per unit of breeding input.

## Conclusions

In summary, across variable rainfall conditions over two seasons, there was a strong correlation between FCR severity as measured using qPCR at maturity and yield of winter cereals grown in inoculated and non-inoculated treatments under field conditions. By using a multivariate analysis, we have presented a method to increase the scale and reliability of selection for genotypes that are more yield responsive in the presence of disease, which could be used to improve selection within wheat and barley breeding programs. This study demonstrated using commercial genotypes that superior lines can be selected under field conditions. For a further study using a crossing population and selection under field conditions is required to validate gains that could be achieved by implementing this strategy within a breeding program. Our use of qPCR to quantify FCR infection severity in this study allowed a higher degree of resolution between genotypes than traditional visual stem basal browning assessments which improved partitioning of tolerance from partial resistance. This method offered new insights into FCR partial resistance and its relative importance to tolerance in bread wheat and barley along with a more robust, less subjective and lower cost technique to potentially select for both important traits within breeding programs.

## Supplementary Information

The online version contains supplementary material.

## Author contribution statement

AM conceived of the study and coordinated its design and execution. AM and BB conducted the experiments and collected the data from the experiments. AM, BO and BO analysed the data. AM wrote the draft. AM, NY, BO, BB, SS and BO reviewed and wrote the manuscript. All authors read and approved the final manuscript.

## Acknowledgments

The Authors thank Michael McCaig and Tony Goldthorpe for their expert technical contributions. This study was conducted as a co-investment between the NSW Department of Primary Industries and the Grain Research and Development Corporation, under project DAN00175.

## Data availability

All data, models, or code generated or used during the study are available from the corresponding author by request.

## Compliance with ethical standards

Conflict of Interest: The authors declare that the research was conducted in the absence of any commercial or financial relationships that could be construed as a potential conflict of interest.

## Notes

### Competing Interest Statement

The authors have declared no competing interest.

