## Supplementary Table 1 for "Improved breeding for *Fusarium pseudograminarium* (Fusarium crown rot) using qPCR measurement of infection in multi-species winter cereal experiments"

Supplementary Table 1. Best linear unbiased predictors (BLUPs) of genotype effects from the multivariate analysis for, grain yield (t/ha) in inoculated and non-inoculated treatments, FCR severity is qPCR of *F. pseudograminarium* in Log_10_ pgDNA/gram, back-transformed FCR severity is pgDNA/gram, the derived trait of yield retention (t/ha) (regression of residuals), the derived trait of tolerance (t/ha) (regression residual), rank position of the genotypes for, inoculated FCR severity, Inoculated yield, retention, tolerance and traditional rating.

|  | Crop | Yield Non-inoculated (t/ha) | Yield inoculated (t/ha) | FCR severity Non-inoculated (Log_10_ pgDNA/gram) | Back transformed non-inoculated CR severity (pgDNA/gram) | FCR severity Inoculated (Log_10_ pgDNA/gram) | Back transformed inoculated CR severity (pgDNA/gram) | Yield Retention | Tolerance | qPCR Inoulated FCR severity rank | Inoculated yld rank | Retention rank | Tolerance rank | Traditional rating |
| --- | --- | --- | --- | --- | --- | --- | --- | --- | --- | --- | --- | --- | --- | --- |
| HINDMARSH | Barley | 3.638 | 3.413 | 0.795 | 6 | 3.533 | 3409 | 0.070 | -0.176 | 1 | 3 | 4 | 24 | S |
| COMPASS | Barley | 3.745 | 3.570 | 0.996 | 10 | 3.671 | 4686 | 0.120 | 0.123 | 2 | 1 | 1 | 5 | S |
| BULOKE | Barley | 3.588 | 3.376 | 0.886 | 8 | 3.686 | 4853 | 0.083 | -0.056 | 3 | 5 | 2 | 17 | SVS |
| LA TROBE | Barley | 3.581 | 3.323 | 0.891 | 8 | 3.705 | 5076 | 0.037 | -0.089 | 4 | 7 | 11 | 21 | MSS |
| COMMANDER | Barley | 3.644 | 3.414 | 0.980 | 10 | 3.751 | 5642 | 0.065 | 0.049 | 5 | 2 | 5 | 8 | MSS |
| SPARTACUS | Barley | 3.592 | 3.291 | 0.957 | 9 | 3.771 | 5902 | -0.005 | -0.053 | 6 | 9 | 16 | 16 | S |
| ROSALIND | Barley | 3.552 | 3.261 | 0.994 | 10 | 3.861 | 7266 | 0.004 | 0.008 | 9 | 13 | 13 | 10 | MSS |
| BASS | Barley | 3.557 | 3.315 | 1.215 | 16 | 4.120 | 13170 | 0.053 | 0.328 | 25 | 8 | 9 | 1 | S |
| EMU ROCK | Bread wheat | 3.521 | 3.288 | 0.941 | 9 | 3.823 | 6645 | 0.062 | -0.004 | 7 | 10 | 6 | 13 | MSS |
| LANCER | Bread wheat | 3.413 | 3.118 | 0.859 | 7 | 3.829 | 6740 | 0.001 | -0.168 | 8 | 17 | 14 | 23 | MSS |
| SUNTOP | Bread wheat | 3.454 | 3.176 | 0.926 | 8 | 3.870 | 7405 | 0.018 | -0.068 | 10 | 14 | 12 | 19 | MSS |
| MERLIN | Bread wheat | 3.513 | 3.277 | 0.986 | 10 | 3.884 | 7663 | 0.060 | 0.049 | 11 | 11 | 7 | 9 | MSS |
| SCEPTER | Bread wheat | 3.512 | 3.262 | 1.004 | 10 | 3.911 | 8151 | 0.046 | 0.060 | 12 | 12 | 10 | 7 | S |
| TROJAN | Bread wheat | 3.603 | 3.385 | 1.083 | 12 | 3.915 | 8215 | 0.077 | 0.187 | 13 | 4 | 3 | 2 | MS |
| BECKOM | Bread wheat | 3.581 | 3.344 | 1.079 | 12 | 3.933 | 8569 | 0.057 | 0.165 | 14 | 6 | 8 | 3 | S |
| CORACK | Bread wheat | 3.464 | 3.165 | 0.999 | 10 | 3.952 | 8962 | -0.002 | 0.006 | 15 | 15 | 15 | 11 | S |
| BOLAC | Bread wheat | 3.481 | 3.116 | 1.044 | 11 | 3.985 | 9665 | -0.069 | -0.010 | 16 | 19 | 20 | 15 | S |
| DS PASCAL | Bread wheat | 3.428 | 3.104 | 1.008 | 10 | 3.997 | 9934 | -0.027 | -0.009 | 17 | 20 | 17 | 14 | S |
| CONDO | Bread wheat | 3.401 | 3.012 | 1.009 | 10 | 4.024 | 10574 | -0.093 | -0.073 | 18 | 22 | 22 | 20 | S |
| FLANKER | Bread wheat | 3.397 | 3.066 | 1.017 | 10 | 4.041 | 11000 | -0.035 | -0.002 | 19 | 21 | 18 | 12 | S |
| EGA_GREGORY | Bread wheat | 3.412 | 3.000 | 1.033 | 11 | 4.045 | 11100 | -0.116 | -0.064 | 20 | 23 | 23 | 18 | S |
| DS DARWIN | Bread wheat | 3.331 | 2.873 | 0.991 | 10 | 4.070 | 11738 | -0.162 | -0.166 | 22 | 24 | 25 | 22 | S |
| PHANTOM | Bread wheat | 3.452 | 3.116 | 1.094 | 12 | 4.076 | 11923 | -0.040 | 0.084 | 23 | 18 | 19 | 6 | MSS |
| WAAGAN | Bread wheat | 3.497 | 3.126 | 1.163 | 15 | 4.115 | 13017 | -0.076 | 0.133 | 24 | 16 | 21 | 4 | S |
| BELLAROI | Durum wheat | 3.228 | 2.803 | 0.896 | 8 | 4.052 | 11262 | -0.128 | -0.255 | 21 | 25 | 24 | 25 | VS |
| Standard error |  | 0.183 | 0.230 | 0.136 |  | 0.170 |  |  |  |  |  |  |  |  |
